# Genome Report: Pseudomolecule-scale genome assemblies of *Drepanocaryum sewerzowii* and *Marmoritis complanata*

**DOI:** 10.1101/2024.04.23.590777

**Authors:** Samuel J. Smit, Caragh Whitehead, Sally R. James, Daniel C. Jeffares, Grant Godden, Deli Peng, Hang Sun, Benjamin R. Lichman

**Affiliations:** Centre for Novel Agricultural Products, Department of Biology, University of York, York, UK; Bioscience Technology Facility, Department of Biology, University of York, York, UK; York Biomedical Research Institute, Department of Biology, University of York, York, UK; Florida Museum of Natural History, University of Florida, Gainesville, FL, USA; School of Life Science, Yunnan Normal University, Kunming, Yunnan, China; Key Laboratory of Yunnan for Biomass Energy and Biotechnology of Environment, Yunnan Normal University, Kunming, China; Key Laboratory for Plant Diversity and Biogeography of East Asia/Yunnan Key Laboratory for Integrative Conservation of Plant Species with Extremely Small Populations, Kunming Institute of Botany, Chinese Academy of Sciences, Kunming, China

**Keywords:** Lamiaceae, Nepetinae, chromosome-level assembly, Nanopore sequencing, Hi-C sequencing

## Abstract

The Nepetoideae, a subfamily of Lamiaceae (mint family), is rich in aromatic plants, many of which are sought after for their use as flavours and fragrances or for their medicinal properties. Here we present genome assemblies for two species in Nepetiodeae: *Drepanocaruym sewerzowii* and *Marmoritis complanata*. Both assemblies were generated using Oxford Nanopore Q20+ reads with contigs anchored to nine pseudomolecules that resulted in 335 Mb and 305 Mb assemblies, respectively, and BUSCO scores above 95% for both the assembly and annotation. We furthermore provide a species tree for the Lamiaceae using only genome derived gene models, complementing existing transcriptome and marker-based phylogenies.

## Introduction

The mint family (Lamiaceae) is the sixth largest plant family with a number of species regarded as important for medicinal, aromatic and ornamental properties (Harley, R, M *et al*. 2004; Zhao *et al*. 2021; Rose *et al*. 2022). Within the Lamiaceae, species from the Nepetoideae are renowned for the accumulation of terpenoids, with tissues used for the extraction of essential oils or as traditional herbal medicines (Wink 2003; Frezza *et al*. 2019). The clade includes widely recognised aromatic species such as mint, lavender, lemon balm and catnip; the volatile terpenoids produced by these plants are responsible for their characteristic fragrances. The ethnobotanical and commercial relevance of this plant family has resulted in considerable scientific interest, including genome assemblies for 36 species at the time of writing (“Published Plant Genomes”).

Here we present the genome assemblies for two Nepetoideae species, namely *Drepanocaryum sewerzowii* (Regel) Pojark. and *Marmoritis complanata* (Dunn) A.L.Budantzev. *M. complanata* is endemic to the subnival band of the Himalaya-Hengduan Mountains, a unique arctic-alpine region recognised as a biodiversity hotspot (Myers *et al*. 2000; Sun *et al*. 2017). This unique habitat necessitates careful control of seed germination to ensure survival (Peng *et al*. 2018). *M. complanata* and other species of the genus are also used as traditional herbal medicines to treat a variety of ailments that include digestive, reproductive, musculoskeletal and skin disorders (Zaman *et al*. 2022). *D. sewerzowii* is native to a region that ranges from Iran to Central Asia and Pakistan and is the sole representative of this genus (Serpooshan *et al*. 2018).

These two species are part of the Nepetinae, a subtribe of the mint family (Lamiaceae, subfamily Nepetoideae, tribe Mentheae) that consists of 375 species and 9-12 genera of which *Nepeta* L. is considered the type genus encompassing 200-300 species. Other genera in this subfamily include *Dracocephalum* L., *Hymenocrater* Fisch. & C.A. Mey., *Lophanthus* Adans., *Agastache* Clayton ex Gronov. and *Schizonepeta* (Benth.) Briq. (Serpooshan *et al*. 2018; Rose *et al*. 2023). The phylogenetic relationship of *M. complanata* and *D. sewerzowii* relative to *N. cataria* L., *N. racemosa* Lam., *A. rugosa* (Fisch. & C.A. Mey.) Kuntze and *S. tenuifolia* (Benth.) Briq. is what prompted our efforts to assemble these genomes. We have been exploring the evolutionary, genomic and enzymatic innovations of monoterpenoid biosynthesis in these species (Lichman *et al*. 2019, 2020; Hernández Lozada *et al*. 2022; Liu *et al*. 2023). However, the available genomic resources provide limited taxonomic coverage. The genome assemblies presented here will allow us to further explore the evolutionary innovations that have impacted terpenoid biosynthesis in the mint family.

## Methods and Materials

### Plant growth conditions

*D. sewerzowii* seeds were obtained from the Millennium Seed Bank at the Royal Botanic Gardens, Kew (serial no. 0694027). *M. complanata* seeds were collected from Puyong Pass Shangri-la County, Yunnan Province, SW China (99°55′E, 28°24′N), 4620 m a.s.l (Peng *et al*. 2018). Seeds were germinated on 1% water agar in a growth room set to 16 h day length, temperature of 20 (±2) °C, relative humidity of 60% (±10%) and a NS12 light spectrum at 120 µmol m^-2^s^-1^ PPFD using Valoya L28 LED lights (Helsinki, Finland). Once a radical emerged, the seedlings were transferred to 7 cm square pots containing Levington Advance Seed and Modular FS2 (ICL Professional Horticulture) seedling soil that was pre-treated with Calypso (Bayer). Once established, a single individual was selected and maintained as a clonal population by propagation using cuttings.

### Genome size estimation by FCM

Genome size estimations were performed through flow cytometry (FCM) using the method of (Dolezel *et al*. 2007). Briefly, the LB01 buffer was used together with *N. cataria* tissue to prepare a reference standard with a previously reported genome size (Mint Evolutionary Genomics Consortium 2018). A CytoFLEX LX (Beckman Coulter) flow cytometer with a 561 nm excitation laser, 610/20 emission filter and a flow rate of 30 µL/min was used. The threshold was set to 488 nm forward scatter to exclude instrument noise and background signal from the buffer.

### Nucleic acid isolation

#### HMW DNA isolation and sequencing

High molecular weight (HMW) DNA was extracted in duplicate from ∼1 g of young leaf tissue using the Nucleobond HMW DNA Extraction kit (Macherey-Nagel, Germany). HMW DNA purity and concentration was assessed by Nanodrop and Qubit, whereafter the extractions were combined. Small fragment DNA elimination was performed with the Circulomics short read eliminator kit (PacBio). Briefly, an equal volume of SRE reagent was added to the sample, and this was centrifuged for 1 h at 12,000 x *g*. The pellet was washed with 70% ethanol before resuspending in TE buffer with low EDTA. DNA quality and quantity was assessed with a nanodrop spectrophotometer (Thermo Fischer Scientific), Agilent Tapestation (running genomic DNA screentape) and Qubit fluorimeter (Invitrogen). Sequencing was performed with the ligation sequencing kit SQK-LSK114 (Oxford Nanopore Technologies), as per the manufacturer’s guidelines, with limited modifications; namely extending the reaction times for end preparation to 30 min at each temperature, and extending adapter ligation steps to an hour). Sequencing was performed on a single promethION FLO-PRO114 flowcell (Oxford Nanopore Technologies) per species, with nuclease flush and sample reload steps performed every 24 h through the run time. For *M. complanata* two additional runs using the SQK-LSK112 ligation sequencing kit (Oxford Nanopore Technologies) and FLO-MIN112 minION flowcells (Oxford Nanopore Technologies) were performed.

Super accuracy base calling was performed using guppy (Oxford Nanopore Technologies) version 6.1.5 for *D. sewerzowii* and version 6.3.9 for *M. complanata*. Read length and quality was assessed using Nanoplot (De Coster and Rademakers 2023). *D. sewerzowii* reads were filtered for a 10 kb minimum length using Nanofilt (De Coster *et al*. 2018). For *M. complanata* we combined all reads from the promethION and minION runs and then filtered using Nanofilt (De Coster *et al*. 2018) with a 3 kb length and Q15 quality cutoff .

#### Genomic DNA isolation and Illumina sequencing

Genomic DNA (gDNA) was extracted from 100 mg of young leaf tissue, in duplicate, using a CTAB extraction method (Doyle and Doyle 1990) and treated with RNAse A. Removal of RNA was confirmed through gel electrophoresis followed by gDNA quality and quantity assessment with a nanodrop spectrophotometer and a Qubit fluorometer (Invitrogen). A total of 508 ng and 752 ng of gDNA for *D. sewerzowii* and *M. complanata*, respectively, was sent for library preparation and paired-end Illumina sequencing with Novogene (Cambridge, UK).

#### RNA isolation and sequencing

RNA was extracted from 80-100 mg of tissue with the Direct-Zol RNA extraction kit (Zymo Research, CA, USA) as per the manufacturer guidelines. For *D. sewerzowii* young and mature leaves, closed and open flowers and stems were used. For *M. complanata* root, young and mature leaf and stem tissues were used. RNA quality was assessed with an Agilent bioanalyzer. Library preparation and paired-end Illumina sequencing was performed by Novogene (Cambridge, UK).

#### Hi-C sequencing

Freshly harvested young leaf leaf tissue was fixed in 1% formaldehyde and washed as per the Phase Genomics (Seattle, WA, USA) sample preparation protocol. Following fixation, the tissue was flash frozen in liquid nitrogen and homogenised using a tissue lyser. The Hi-C libraries were prepared and sequenced by Phase Genomics (Seattle, WA, USA).

### Genome Assembly

Filtered nanopore reads for the respective genomes were used for assembly and error correction. Both species were first assembled using Flye (Lin *et al*. 2016; Kolmogorov *et al*. 2019) (--iterations 0 and --nano-hq flags). *M. complanata* was also assembled with NECAT (Chen *et al*. 2021) using the default configuration file settings. Our error correction pipeline entailed polishing with long reads by two rounds of RACON (Vaser *et al*. 2017), with reads mapped using minimap2 (Li 2018), followed by two rounds of MEDAKA (*medaka: Sequence correction provided by ONT Research* 2018) polishing. Short reads were mapped using bwa-mem (Li 2013) and duplicate reads marked using Picard (*Picard toolkit* 2019) prior to two iterative rounds of polishing with Pilon (Walker *et al*. 2014).

The *M. complanata* Flye and NECAT assemblies were merged with Quickmerge (Chakraborty *et al*. 2016; Solares *et al*. 2018) due to the low N50 scores. The overlap cutoff (-c flag) was five and the length cutoff (-l) was 100,000 with the NECAT assembly used as the query. The NECAT-Flye merged assembly underwent another two rounds of short read error correction using Pilon. For *M. complanata* we purged the merged assembly of haplotigs prior to HiC scaffolding while *D. serwerzowii* was purged after HiC scaffolding. Haplotig purging was performed using the purge haplotigs pipeline (Roach *et al*. 2018). Contigs were scaffolded into pseudomolecules by Phase Genomics (Seattle, WA, USA) using the Proximo Genome Scaffolding Platform. Contiguity and completeness was assessed throughout the assembly pipeline using BUSCO (Benchmarking for University Single Copy Orthologs) v5.4.2 with the embryophyta_odb10 dataset (Manni *et al*. 2021).

### Genome annotation

Repeats and transposable elements were annotated using the Earl Grey v3.2 (Baril *et al*. 2023, 2024) pipeline with default settings followed by softmasking of the repeats using the maskfasta function of bedtools. The BRAKER3 pipeline (v3.0.6) (Stanke *et al*. 2006, 2008; Gotoh 2008; Iwata and Gotoh 2012; Buchfink *et al*. 2015; Hoff *et al*. 2016, 2019; Kovaka *et al*. 2019; Pertea and Pertea 2020; Brůna *et al*. 2021; Bruna *et al*. 2024) was used to predict gene models using mRNA and protein evidence. For protein evidence we generated a representative database from 52 Mint species (48 Lamiaceae and four from Lamiales families) using the transcriptomes from (Mint Evolutionary Genomics Consortium 2018). MMseqs2 (Steinegger and Söding 2017) was used to remove identical sequences from the database. For mRNA evidence we aligned RNAseq reads from the different tissues using STAR (Dobin *et al*. 2013) with default settings and the “--outSAMstrandField intronMotif” flag. The respective bam outputs were merged using samtools (Danecek *et al*. 2021) and used as input for BRAKER3. The BRAKER annotation output was reformatted to GFF3 using AGAT (Dainat *et al*. 2023) followed by extraction and translation of the longest open-reading for each predicted coding sequence. Annotation completeness was assessed using BUSCO (Manni *et al*. 2021) in protein mode with the embryophyta_odb10 dataset.

### Species tree and macrosynteny analysis

Markerminer (Chamala *et al*. 2015) was used to identify single-copy genes using predicted coding genes from representative Lamiaceae genomes (Sup. Table 1) and *Paulownia fortunei* (Seem.) Hemsl. as an outgroup. Genes present in 26 of the 27 species were included. The MAFFT alignments generated as part of the Markerminer pipeline were trimmed for gaps using using the gappyout algorithm of trimAl v1.4.1 (Capella-Gutiérrez *et al*. 2009) and concatenated into a supermatrix with partitions using the catfasta2phyml script (https://github.com/nylander/catfasta2phyml). A species-tree was inferred by maximum likelihood with partition models (Chernomor *et al*. 2016) using IQ-TREE 2 (Minh *et al*. 2020) with ModelFinder (Kalyaanamoorthy *et al*. 2017), ultrafast bootstraps (UFBoot2, X1000) (Hoang *et al*. 2018), and SH-aLRT supports (X1000) (Guindon *et al*. 2010). In addition, a species tree using protein sequences was inferred using the STAG (Species Tree inference from All Genes) method of Orthofinder (Emms and Kelly 2015, 2017, 2018, 2019). Pairwise macrosynteny analyses were performed against *A. rugosa (Park et al. 2023)* and *S. tenuifolia (Liu et al. 2023)* using the JCVI (Tang *et al*. 2015) implementation of MCScan (Tang *et al*. 2008). MCScan orthologs were identified in full mode with predicted protein sequences and default settings.

**Table 1.** Assembly and annotation metrics.

|  | *D. sewerzowii* | *M. complanata* |
| --- | --- | --- |
| **Assembly Statistics** |  |  |
| Assembly size (Mb) | 332.85 | 305.55 |
| Number of pseudomolecules | 9 | 9 |
| N50 (Mb) | 35.13 | 27.69 |
| L50 | 5 | 5 |
| L90 | 9 | 28 |
| GC% | 38.56 | 37.45 |
| Number of Ns | 2600 | 18 800 |
| **Annotation Statistics** |  |  |
| Assembly BUSCO*<br>*n*=1440 | C: 99.0 %<br>S: 96.3 %<br>D: 2.7 % | C: 95.7 %<br>S: 85.7 %<br>D: 10.0 % |
| Annotation BUSCO*<br>*n*=1440 | C: 95.0 %<br>S: 92.1 %<br>D: 2.9 % | C: 95.3 %<br>S: 86.1 %<br>D: 9.2 % |
| Predicted coding genes | 24 221 | 25 080 |
| Predicted proteins | 26 989 | 28 384 |
| Percentage repeats | Total: 62%<br>DNA: 2.38 %<br>LINE: 0.53%<br>LTR: 40.61% | Total: 53%<br>DNA: 4.43%<br>LINE: 2.92%<br>LTR: 25.83% |

\* Complete ( C), Single (S), Duplicated (D)

### Expression analysis

RNAseq read alignments were evaluated with STAR (Dobin *et al*. 2013) and assed with qualimap (García-Alcalde *et al*. 2012; Okonechnikov *et al*. 2016). Qualimap reports were aggregated with MultiQC (Ewels *et al*. 2016). Expression counts as transcripts per million (TPM) were generated using Salmon (Patro *et al*. 2017). The transcript index for Salmon was generated using the full set of predicted coding sequences from BRAKER3.

## Results and Discussion

### Chromosome level assemblies

We sequenced the genomes for *D. sewerzowii* and *M. complanata* using Oxford Nanopore long reads and Proximo HiC scaffolding (Phase Genomics) resulting in two chromosome-level assemblies. A total of 99.24 Gb of super accurate nanopore reads were generated for *D. sewerzowii* with 80 Gb of reads being greater than 10 kb at a mean read quality (Q-score) of 16.6. The size filtered reads provided 242X coverage at an estimated genome size of 330 Mb, as determined through FCM (Supl. Fig. 1). The initial Flye assembly resulted in 472 contigs, an N50 of 17 Mb, a total assembly length of 333.75 Mb and a BUSCO score of 98.7%. Polishing with long and short reads reduced the number of contigs to 134 and assembly size to 332.85 Mb while maintaining a N50 of 17 Mb. The BUSCO score increased slightly to 98.8% after polishing. HiC scaffolding orientated the assembly to nine pseudomolecules (Supl. Fig 2A), which is in agreement with the chromosome counts reported by Bordbar (2023). The nine pseudomolecules contained 97.6% of the contigs, representing 324.87 Mb of the total assembly at a N50 of 35.2 Mb and L50 of 5 (Table 1).

**Figure 1.**
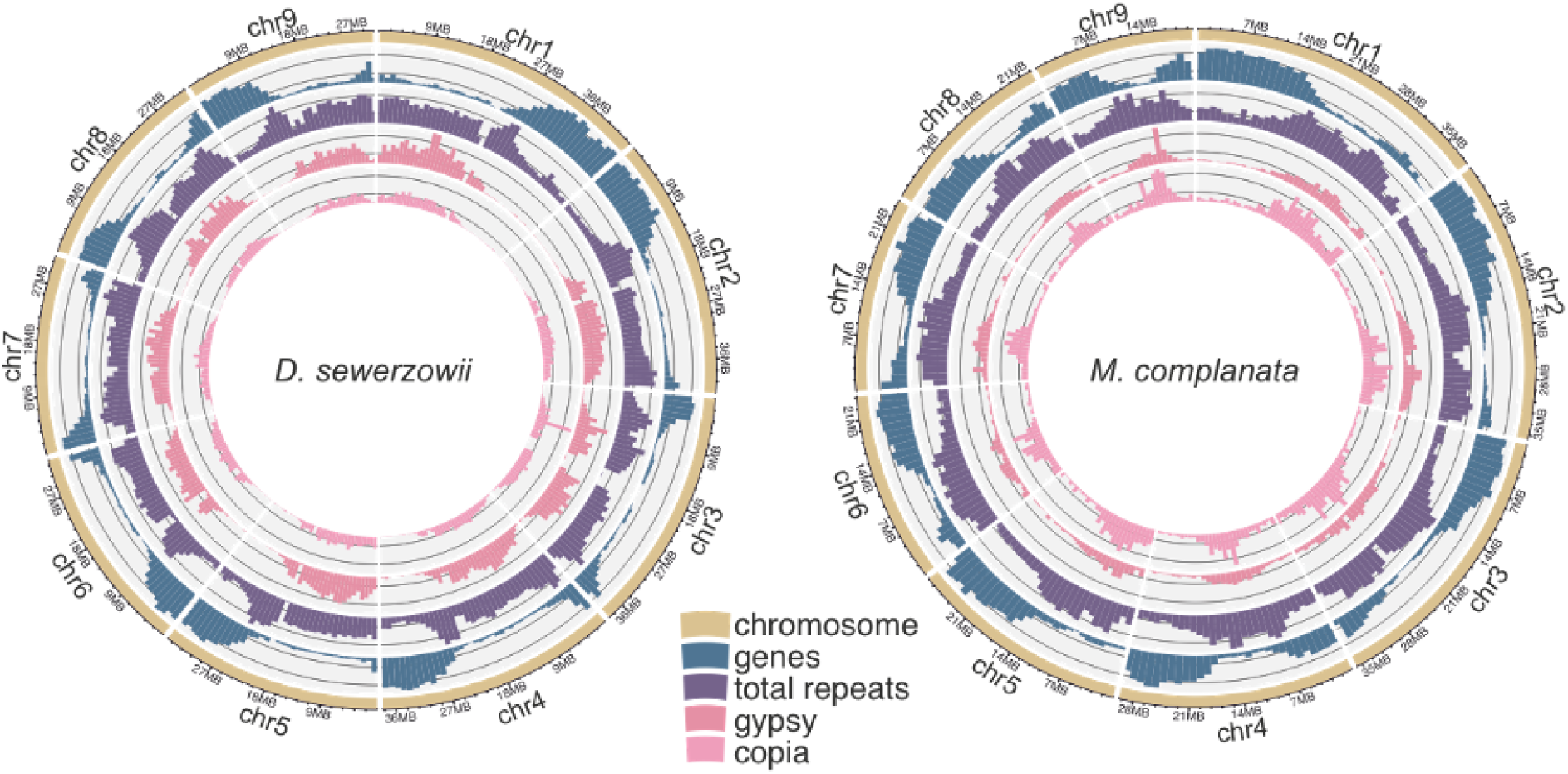
Circos plots for the genome assemblies of *D. sewerzowii* and *M. complanata* depicting density (1 Mb bins) of genes, total repeats, gypsy and copia elements along the 9 pseudomolecules.

**Figure 2.**
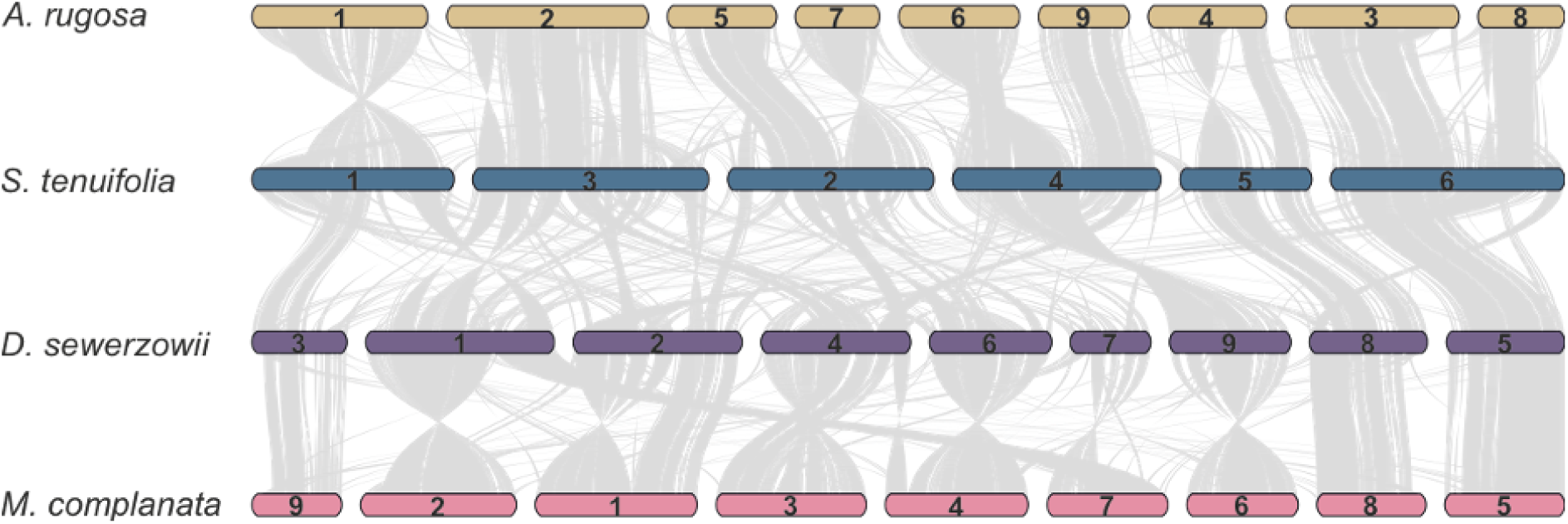
Pairwise macrosynteny analysis of the assembled genomes relative to closely related species with chromosome level assemblies. Conserved collinear blocks are linked by the grey lines.

The *M. complanata* genome size was estimated at 337 Mb using FCM (Supl. Fig. 1). We obtained 59.3 Gb of reads after length and quality filtering, providing 176X coverage with a mean Q-score of 18. We tried various different read filtering cutoffs for both length and quality with all attempts using Flye failing to reach a N50 greater than ∼335 kb. After polishing the best Flye assembly was 420 Mb in size with a N50 of 335 kb, 3001 contigs and a BUSCO score of 98.5%, of which 17.1% were duplicated. NECAT resulted in a more contiguous genome assembly of 457 Mb with a N50 of 1 Mb, 869 contigs and 98.6% BUSCO, of which 36.7% were duplicated. The inflated genome size and high number of duplicate BUSCO genes suggested that the fragmented assemblies contained a high number of haplotigs (contigs of a single haplotype), that would artificially inflate genome size.

In an attempt to increase the continuity of the assembly (N50 score) we merged the Flye and NECAT assemblies. The NECAT assembly had fewer contigs and greater N50 and was therefore selected to be the query genome with the Flye assembly used to improve the query genome. We evaluated the impact of haplotig purging before and after merging. Each assembly was purged of haplotigs prior to merging and compared to a merged assembly that was purged as the final step. In each iteration we polished twice with short reads after merging. The merging increased the N50 to 3 Mb regardless of when we purged the haplotigs. The timing of the purging step had a large impact on the number of contigs together with a minor impact on the duplicated BUSCOs. Merging, polishing and then purging the haplotigs resulted in the most contiguous assembly (305.6 Mb) with the fewest number of contigs (338) and a BUSCO score of 95.6 %. HiC scaffolding assembled the contigs into nine pseudomolecules (Supl. Fig 2B), which is in agreement with karyotype information (Sun 2016), totaling 258 Mb (85% of the total assembly). The pseudomolecules had a BUSCO score of 91.3% with the total assembly having a BUSCO of 95.7% (Table 1).

Pseudomolecule termini were manually inspected for presence of the TTTAGGG telomeric repeat. Seven of the *D. sewerzowii* pseudomolecules contained this repeat on at least one end with chr. 4 and 7 having it on both ends. For *M. complanata* we found this repeat on six pseudomolecules with chr. 7 and 8 having it on both ends. The presence of this repeat on both ends indicates a telomere to telomere assembly for these chromosomes.

### Repeat and genome annotations

Repeat annotation revealed that 62% of the *D. sewerzowii* genome and 53% of the *M. complanata* genome are repeats (Table 1). The largest portion of the repeats were long terminal repeats (LTR), occupying 40.6% and 25.8% of the respective genomes. Subsequent to repeat masking our gene annotation, using *ab initio*, protein and mRNA predictions, resulted in 24,221 and 25,080 gene regions that encode for 26,989 and 28,384 proteins for the respective genomes. BUSCO analysis of the primary isoforms was 95% for both genomes. Gene and repeat density showed an inverse relationship along the chromosomes (Figure 1).

The RNAseq data we produced found evidence for expression of the majority of genes. RNAseq reads mapped to gene models showed that 85% (22,794/26,815) of the genes were expressed in at least one tissue type for *D. sewerzowii* and 88% (24,919/28,384) of the genes in *M. complanata*. Expression matrices as transcripts per million (TPM) are available in Supplementary Tables 3 and 4.

### Pairwise macrosynteny and species tree

We compared our assemblies to the closest relatives with pseudomolecule assemblies, namely *A. rugosa* (Park *et al*. 2023) and *S. tenuifolia* (Liu *et al*. 2023) (Figure 2). *A. rugosa* has a 9 chromosome assembly with macrosynteny revealing that the overall genome structures for both *D. sewerzowii* and *M. complanata* are similar to this species. Macrosynteny revealed a number of chromosome fusion events in *S. tenuifolia* relative to the other three genomes. Although only 85% of the *M. complantum* contigs were anchored to pseudomolecules the overall structure of the chromosomes (relative to *A. rugosa* and *D. sewerzowii*) and presence of large syntenic blocks indicate a reasonably complete assembly.

The phylogenetic relationships of the Lamiaceae have been reported using plastid, nuclear and transcriptome approaches. The species trees presented in Figure 3 used genome derived gene models for phylogenomic inference, complementing existing species trees (Serpooshan *et al*. 2018; Mint Evolutionary Genomics Consortium 2018; Rose *et al*. 2022, 2023). The STAG species-tree used multi-copy gene families (i.e. orthogroups) predicted by Orthofinder using protein sequences. The consensus tree in Fig 3A shows internal bipartition support for 5,296 orthogroups in which all species are present. The ML tree (Fig. 3B) was inferred from a single-copy gene supermatrix totaling 340,706 nucleotide sites with all but two branches showing above 98% support for both ultrafast bootstraps and SH-aLRT. The two branches indicated by the asterisk were not well supported, bootstrap and SH-aLRT

**Figure 3.**
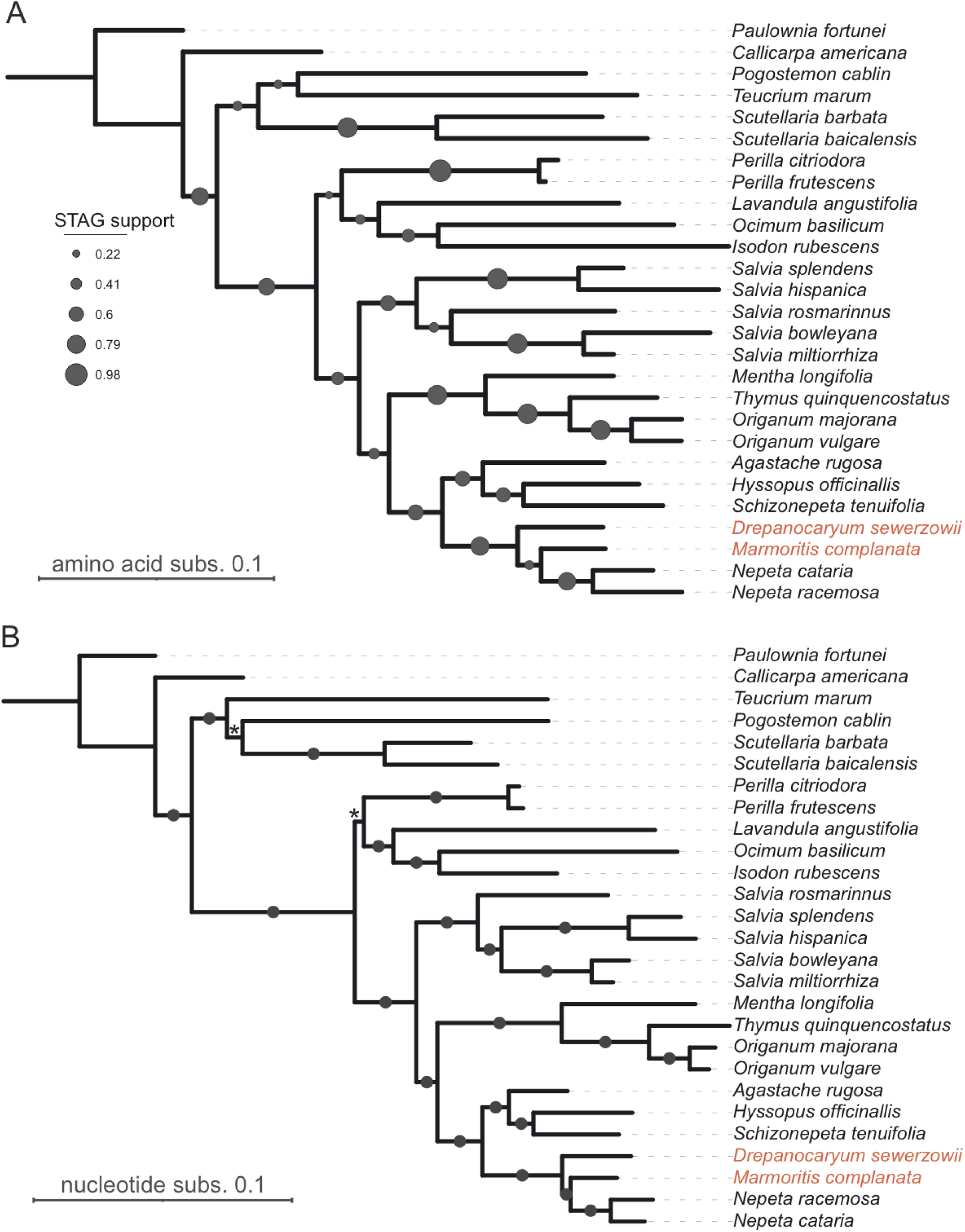
Species-trees inferred with Lamiaceae genome derived gene models. (A) STAG species-tree inferred with Orthofinder protein orthogroups. Support values show the proportion of trees at which the internal bipartitions occur for all species. (B) Maximum-likelihood species tree using single copy nucleotide sequences. Branches with circles are fully supported (>98%) as judged by ultrafast bootstraps and SH-aLRT. Branches indicated by the asterisk are less well supported (<85%).

<85%.

In both the ML and STAG topologies, *D. sewerzowii* is recovered as a sister to a clade that includes *M. complanata* and *Nepeta*, which are sister to each other. While our phylogenomic results corroborate existing hypotheses regarding the close relationships among these genera, our trees are incongruent with previously reported topologies (Supl. Fig. 3). For example, nuclear phylogenetic results by Rose *et al*. (2023) report *D. sewerzowii* as sister to *Nepeta*, which together are sister to the sister lineages *Hymenocrater* and (*Lophanthus* + *Marmoritis*). This contrasts with plastid-based phylogenetic results reported in the same study, which recover *Nepeta* as sister to a clade comprising the sister taxa, *Drepanocaryum* and *Hymenocrater*, and their sister, (*Lophanthus* + *Marmoritis*), and with results by Serrpooshan *et al*. (2018), which recover *D. sewerzowii* as sister to a mixed and partially unresolved clade of *Hymenocrater*, *Lophanthus*, *Marmoritis*, and *Nepeta*. Topological discordances among trees reported in this and previous studies likely reflect differences in taxonomic and molecular sampling, but they also highlight the complexity of resolving intergeneric relationships within Nepetinae. The species-tree presented here (Fig. 3) provides necessary context for comparative genomics, although interpretations should be considered alongside available transcriptome- and marker-based phylogenies until additional Nepetinae genomes and phylogenomic results become available. Nevertheless, the genomes presented here provide a valuable resource to explore the evolutionary trajectories underpinning the remarkable innovations in specialised metabolism within the Lamiaceae.

## Conclusion

Plant genome assemblies are being generated at a remarkable rate, with two-thirds of available plant genome assemblies generated within the last 3 years (Xie *et al*. 2024). Here we present the chromosome-level genome assemblies of *D. sewerzowii* and *M. complanata*, representing the first assemblies from these genera. The gene and repeat annotations, along with expression matrices, present a comprehensive resource for comparative genomics. The species-tree using gene models from available Lamiaceae genome assemblies provides a reference point that celebrates the number of sequenced species. These genome assemblies will allow us to decipher the evolutionary innovations that resulted in the remarkably diverse number of specialised metabolites found in the Lamiaceae.

## Data Availability Statement

The raw reads for whole genome and transcriptome sequencing are available in the National Center for Biotechnology Information Sequence Read Archive BioProject PRJNA1097548 and PRJNA1095452. The genome assembly, annotation files and gene expression abundance datasets are available through Figshare as supplementary data.

## Acknowledgments

We would like to thank Prof. C. Robin Buell for her advice and on assembling mint genomes. The Viking cluster was used during this project, which is a high performance compute facility provided by the University of York. We are grateful for computational support from the University of York, IT Services and the Research IT team. We are grateful to the University of York Horticulture Team for the propagation and care of our plant material. We would like to thank Karen Hobb from the University of York Imaging and Cytometry Laboratory for assistance with FCM.

## Supplementary Data

**Supplementary Figure 1.**
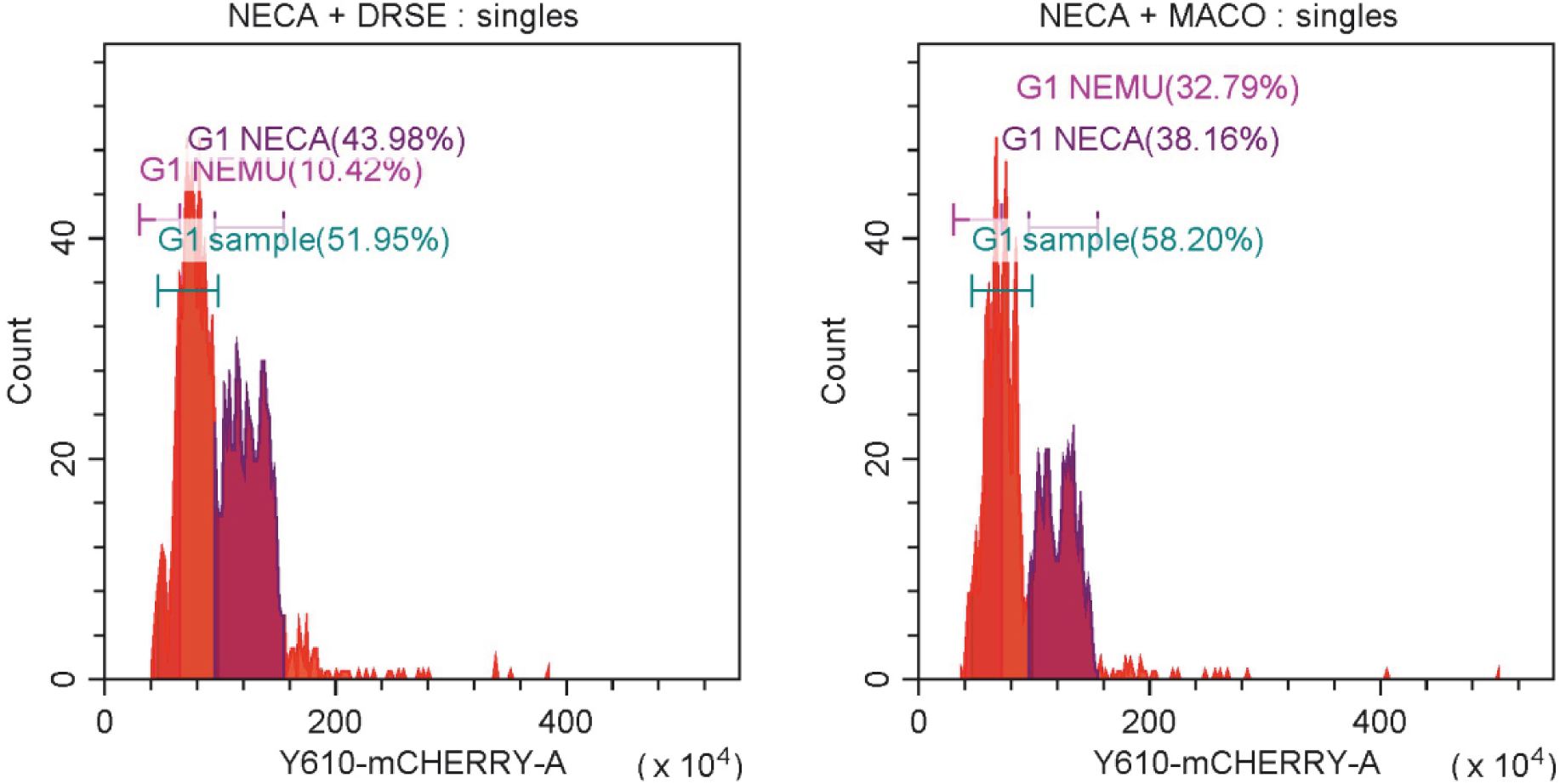
FCM ungated histograms for *D. sewerzowii* (A) and *M. complanata* (B). *N. cataria* tetraploid (G1 NECA) and *N. racemosa* diploid (G1 NEMU) references are shown relative to that of *D. sewerzowii* and *M. complanata*, labelled as G1 sample.

**Supplementary Figure 2.**
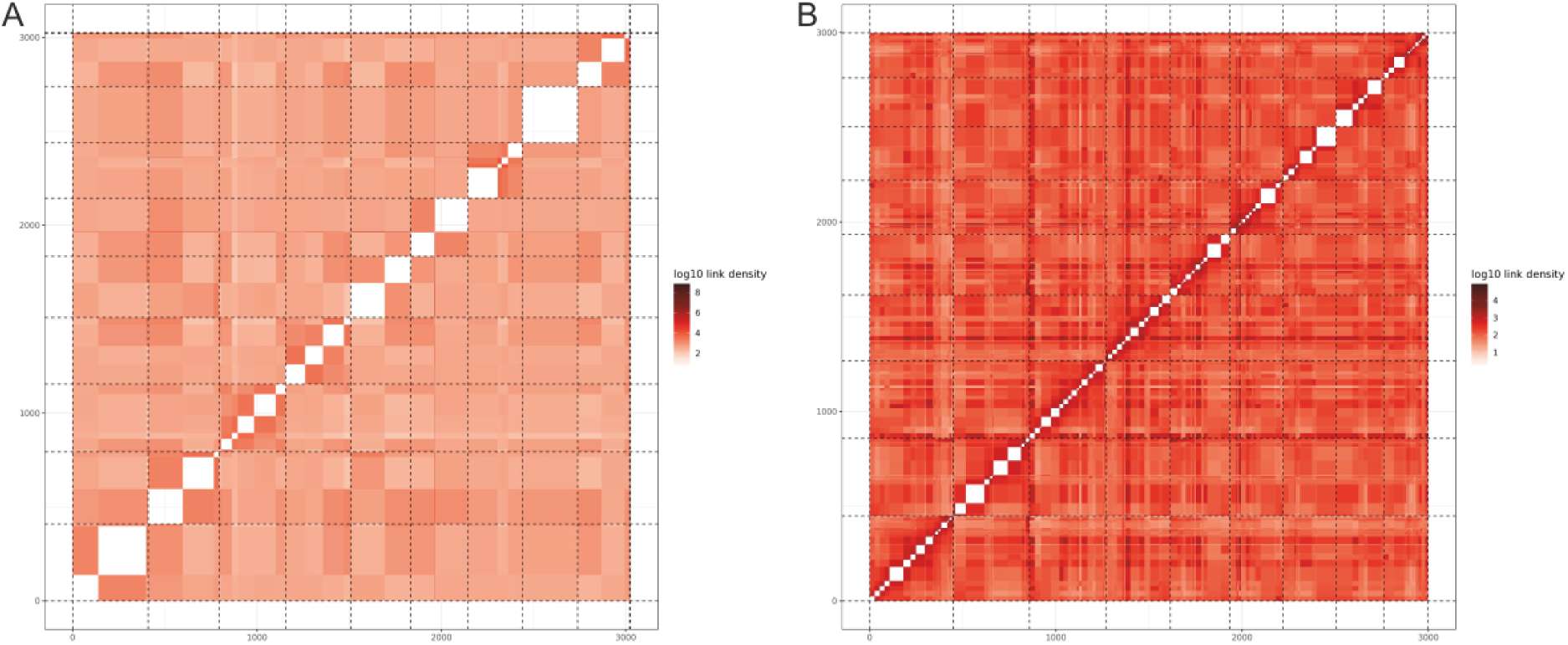
HiC contact maps for *D. sewerzowii* (A) and *M. complanata* (B).

**Supplementary Figure 3.**
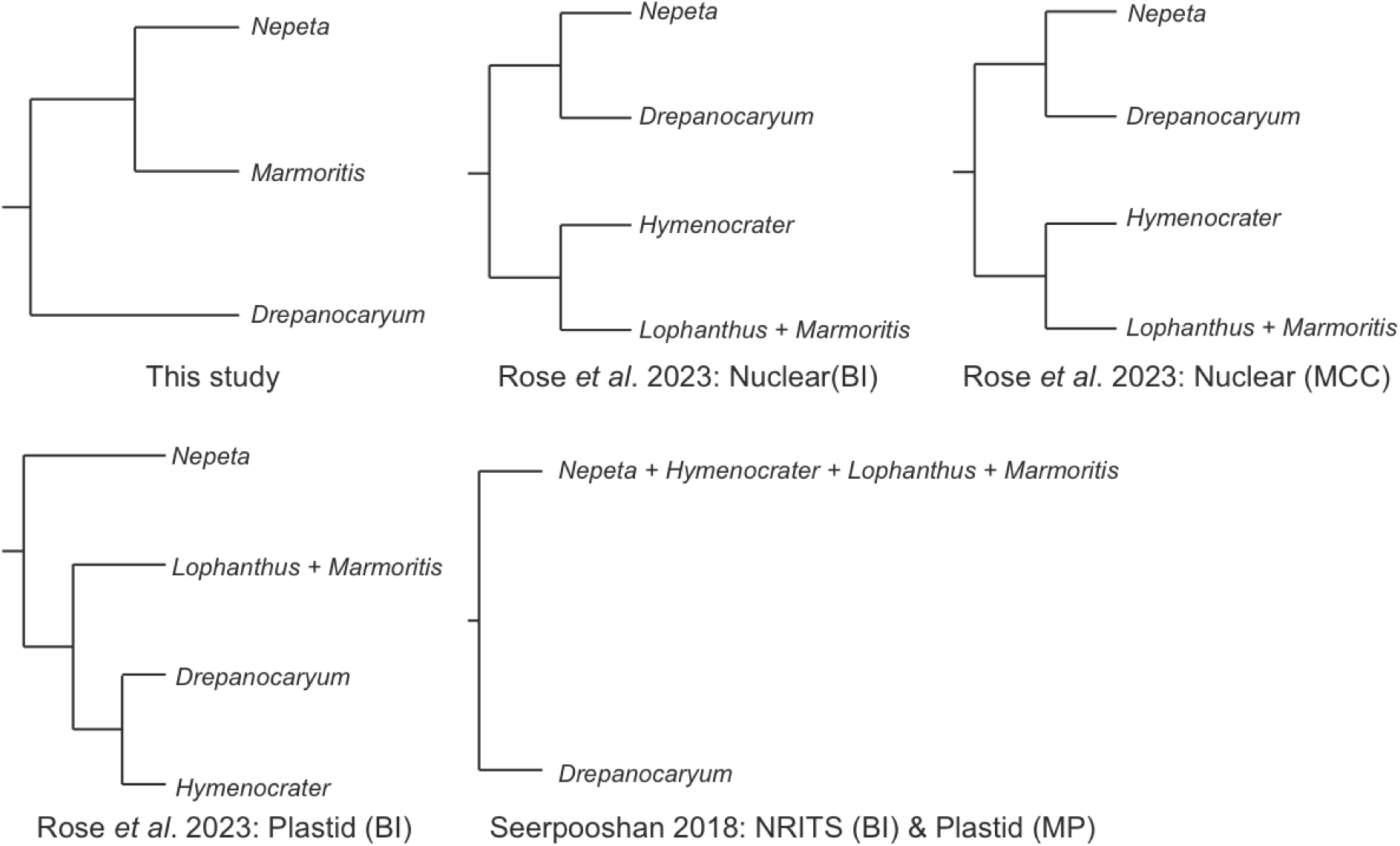
Summary of tree incongruence between the genome derived species tree (this study) and trees reported using Bayesian Inference (BI), maximum clade credibility (MCC) or maximum parsimony (MP) for tree inference with plastid markers, nuclear markers or nuclear ribosomal internal transcribed spacer regions (NRITS).

**Supplementary Table 1.** Genome assemblies used for comparative genomics and phylogenomics.

| Species | Reference |
| --- | --- |
| *Agastache rugosa* (Fisch. & C.A.Mey.) Kuntze | Park *et al.* 2023 |
| *Callicarpa americana* L. | Hamilton *et al.* 2020 |
| *Drepanocaryum sewerzowii* (Regel) Pojark. | This work |
| *Hyssopus officinalis* L. | Lichman *et al.* 2020 |
| *Isodon rubescens* (Hemsl.) H.Hara | Sun *et al.* 2023 |
| *Lavandula angustifolia* Mill. | Hamilton *et al.* 2023 |
| *Marmoritis complanata* (Dunn) A.L.Budantzev | This work |
| *Mentha longifolia* (L.) L. | Vining *et al.* 2022 |
| *Nepeta cataria* L. | Lichman *et al.* 2020 |
| *Nepeta racemosa* Lam. | Lichman *et al.* 2020 |
| *Ocimum basilicum* L. | Bornowski *et al.* 2020 |
| *Origanum majorana* L. | Bornowski *et al.* 2020 |
| *Origanum vulgare* L. | Bornowski *et al.* 2020 |
| *Paulownia fortunei* (Seem.) Hemsl. | Cao *et al.* 2021 |
| *Perilla citriodora* (Makino) Nakai | Zhang *et al.* 2021 |
| *Perilla frutescens* (L.) Britton | Zhang *et al.* 2021 |
| *Pogostemon cablin* (Blanco) Benth. | Shen *et al.* 2022 |
| *Salvia bowleyana* Dunn | Zheng *et al.* 2021 |
| *Salvia hispanica* L. | Wang *et al.* 2022 |
| *Salvia miltiorrhiza* Bunge | Pan *et al.* 2023 |
| *Salvia rosmarinus* Spenn. | Han *et al.* 2023 |
| *Salvia splendens* Sellow ex Nees | Jia *et al.* 2021 |
| *Schizonepeta tenuifolia* (Benth.) Briq. | Liu *et al.* 2023 |
| *Scutellaria baicalensis* Georgi | Xu *et al.* 2020 |
| *Scutellaria barbata* D.Don | Xu *et al.* 2020 |
| *Teucrium marum* L. | Smit *et al.* 2024 |
| *Thymus quinquecostatus* Čelak. | Sun *et al.* 2022 |

